# Integrated population genomic analysis and numerical simulation to estimate larval dispersal of *Acanthaster* cf. *solaris* between Ogasawara and other Japanese regions

**DOI:** 10.1101/2021.03.25.436911

**Authors:** Mizuki Horoiwa, Takashi Nakamura, Hideaki Yuasa, Rei Kajitani, Yosuke Ameda, Tetsuro Sasaki, Hiroki Taninaka, Taisei Kikuchi, Takehisa Yamakita, Atsushi Toyoda, Takehiko Itoh, Nina Yasuda

## Abstract

The estimation of larval dispersal of marine species occurring on an ecological timescale is significant for conservation. In 2018, a semi-population outbreak of crown of thorns starfish, *Acanthaster* cf. *solaris* was observed on a relatively isolated oceanic island, Ogasawara. The aim of this study was to assess whether this population outbreak was caused by large-scale larval recruitment (termed secondary outbreak) from the Kuroshio region. We estimated larval dispersal of the coral predator *A.* cf. *solaris* between the Kuroshio and Ogasawara regions using both population genomic analysis and oceanographic dispersal simulation. Population genomic analysis revealed overall genetically homogenized patterns among Ogasawara and other Japanese populations, suggesting that the origin of the populations in the two regions is the same. In contrast, a simulation of 26-year oceanographic dispersal indicated that larvae are mostly self-seeded in Ogasawara populations and have difficulty reaching Ogasawara from the Kuroshio region within one generation. However, a connectivity matrix produced by the larval dispersal simulation assuming a Markov chain indicated gradual larval dispersal migration from the Kuroshio region to Ogasawara in a stepping-stone manner over multiple years. These results suggest that, while large-scale larval dispersal from an outbreak of the Kuroshio population spreading to the Ogasawara population within one generation is unlikely. This study also highlighted the importance of using both genomic and oceanographic methods to estimate larval dispersal, which provides significant insight into larval dispersal that occurs on ecological and evolutionary timescales.

## 1 Introduction

Many benthic marine invertebrates in coral reef ecosystems exhibit planktonic larval dispersal during their early life history. Such larval dispersal connects different populations and forms a meta-population structure (Shanks, 2009). Thus, for effective conservation and management of benthic marine invertebrates, assessment of larval dispersal is important (Botsford et al., 2009; Almany et al 2009). Because it is challenging to directly track the movement of tiny, numerous larvae in the field, several different indirect methods, including population genomic analysis (e.g. Yasuda et al., 2015; Arndt and Smith, 2002), oceanographic simulation (e.g. Miyake et al., 2009; Storlazzi et al., 2017), plankton-netting in the field (e.g. Yasuda et al., 2015; Suzuki et al., 2016), and drifting buoys (e.g. Fukuda and Hanamura, 1996), have been employed to estimate larval dispersal. Although each method has advantages and disadvantages, the first two methods can be applied to relatively large spatial scales and thus have been used more frequently than the other methods. On the one hand, the data obtained by population genomic analysis reflect the biological processes of larval dispersal and recruitment. However, they not only include larval dispersal occurring on an ecological timescale but also reflect historical gene flow and genetic breaks caused by past climate change and plate tectonics (Benzie, 1999; Crandall et al., 2014). Oceanographic simulation, on the other hand, can capture snapshots of larval dispersal, while even a sophisticated biophysical model cannot accurately simulate larval behavior, recruitment, and survival in the ocean (White et al., 2019). It is therefore important to integrate different methods to estimate larval dispersal (Marko and Hart, 2017), although few studies have attempted to do so (but see Schunter et al., 2011; Alberto et al., 2011; Nakabayashi et al., 2019; Taninaka et al., 2019).

In this study, we targeted the crown-of-thorns starfish, *Acanthaster* cf. *solaris*, to estimate larval dispersal. Chronic outbreaks of the coral predator *A.* cf. *solaris* and its siblings have been the major management issue in coral reef ecosystems in the Indian and Pacific oceans (Baird et al., 1990; Baird et al., 2013). With high fecundity and a larval stage that lasts a few to several weeks (Yamaguchi, 1977; Lucas, 1973), a population outbreak can cause successive population outbreaks, termed secondary outbreaks (reviewed in Birkeland and Lucas, 1990; Pratchett et al., 2014; Yasuda, 2018), via larval dispersal in neighboring regions, especially in the western Pacific region. Assessment of the spatial range of secondary outbreaks of *A.* cf. *solaris* is necessary for coral reef conservation.

Many population genomic and phylogeographic studies of *A.* cf*. solaris* have been conducted using various molecular markers (e.g., Benzie, 1992, 1999; Vogler et al, 2013; Timmers et al., 2011; Yasuda et al., 2009; 2015). These studies have indicated strong gene flow over a long distance along western boundary currents such as the Kuroshio Current region (genetic homogeneity from across the Ryukyu Islands to the temperate Pacific coast of mainland Japan, Yasuda et al., 2009) and the Eastern Australian Current region (the Great Barrier Reef, Nash et al., 1988; Benzie et al., 1999; Vogler et al., 2013; Harrison et al., 2017), while relatively limited gene flow and genetic differentiation were observed among distant Pacific islands (Yasuda et al., 2009; Timmers et al., 2012; Vogler et al., 2013). Overall, the studies found strong gene flow and genetic homogeneity in regions where strong currents are associated, although genetic data strongly reflect historical gene flow that was formed during past climate change, and the spatial extent of secondary outbreaks could be overestimated (Benzie 1999, Yasuda et al. 2009, Yasuda et al. 2015). Larval dispersal simulation studies in the Great Barrier Reef (GBR) have indicated southward larval dispersal along the Eastern Australian Current, which is consistent with historically observed patterns of secondary outbreaks of *A.* cf*. solaris.* (Dight et al., 1990a, b, James and Scandol, 1992; Scandol and James, 1992). Plankton netting survey has also showed evidence of large-scale larval dispersal of *A.* cf. *solaris* in the GBR (Uthicke et al., 2015). While intensive connectivity studies of *A.* cf. *solaris* have been conducted at the Indo-Pacific scale and in some localized regions where secondary outbreaks were suspected (e.g., the Kuroshio region and the GBR), connectivity between such regions and relatively isolated oceanic island regions is yet to be examined.

In 2018, a population outbreak of *A.* cf. *solaris* on Ogasawara Island was reported (Biodiversity Center of Japan, 2019), which was the second population outbreak on Ogasawara since 1979 (Kurata, 1984). Ogasawara Island is an oceanic island situated ca. 1000 km south of the Tokyo urban center and is out of the mainstream of the Kuroshio Current. It is therefore still unknown whether this population outbreak is attributed to a secondary outbreak in the Kuroshio region.

The aim of this study was to estimate larval dispersal between the Kuroshio region and Ogasawra to determine whether the population outbreak in Ogasawara was caused by a secondary outbreak in the Kuroshio region. To meet this aim, we used both a method of population genomic analysis called multiplexed ISSR genotyping by sequencing (MIG-seq) and oceanographic larval dispersal simulation based on the Global HYbrid Coordinate Ocean Model (HYCOM).

## 2 Material and Methods

### 2.1 Sampling and DNA extraction

A total of 187 starfish samples were collected from six Kuroshio regions (Tatsukushi, Miyazaki, Sakurajima, Onna Village, Miyako, and Sekisei Lagoon) and Ogasawara via scuba diving (Fig. 1). The tube feet were preserved in 99.5% ethanol prior to DNA extraction. Genomic DNA extraction was carried out using a modified heat alkaline method (Meeker at al., 2007; Nakabayashi et al., 2019).

**Figure 1.**
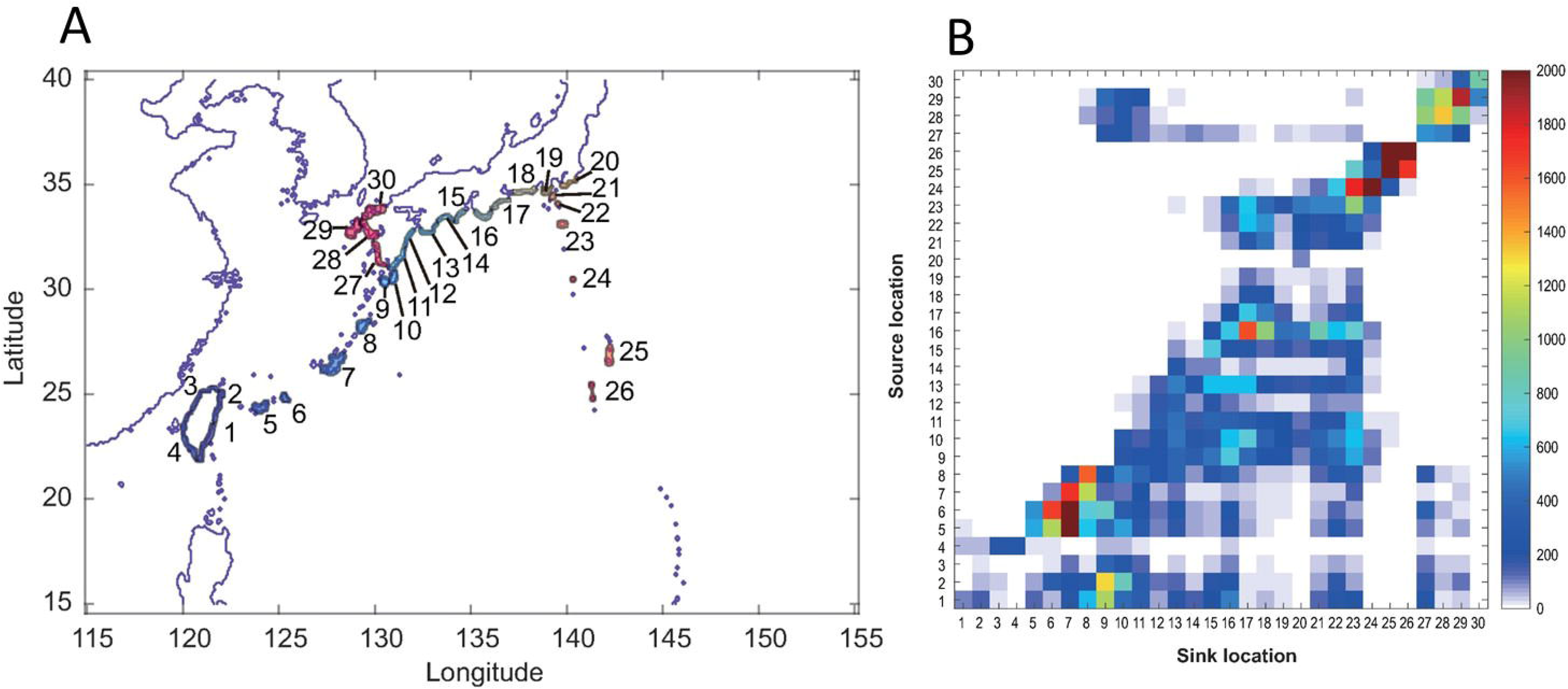
**(A)**: The site numbers for numerical simulation including sampling sites for genomic analysis (5 = Sekisei Lagoon [SKS], 6 = Miyako [MYK], 7 = Onna Village [ONN], 11 = Miyazaki [MYZ], 13 = Tatsukushi [TTK], 25 = Ogasawara [OGS], and 27 = Sakurajima [SKR]). **(B)**: The connectivity matrix of the simulation of 26-year larval dispersal. The sink location is indicated on the x-axis, and the source location is indicated on the y-axis. Colors indicate numbers of particles.

### 2.2 Whole-genome sequencing and nuclear genome assembly as the reference

We obtained a new high-quality whole-genome sequence for mapping analysis. Whole-genome sequencing of the sample collected from Miyazaki (BioSample ID, SAMD00056692) was performed using Sequel, which is a single-molecule real-time sequencer from Pacific Biosciences (PacBio; Menlo Park, CA, USA), according to the manufacturer’s protocol. Previously published whole-genome short-read sequencing data (Wada et al., 2020) were downloaded from the BioProject PRJDB4009 in the ENA/Genbank/DDBJ database. Briefly, we obtained two paired-end (PE) libraries (insert sizes: 300 and 500 bp) and six mate-pair (MP) libraries (insert sizes: 3k, 5k, 8k, 10k, 12k, and 15k).

For the *in silico* procedures in this section, default parameters were used except where otherwise noted. Illumina PE and MP reads were filtered and trimmed using Platanus_trim v1.0.7, (http://platanus.bio.titech.ac.jp/pltanus_trim). We assembled trimmed PE and MP reads using Platanus v1.2.5 (Kajitani et al., 2014) with the commands “assemble,” “scaffold,” and “gap_close,” resulting in the scaffolds.

In addition to the short-read assembly, PacBio long-read assembly was conducted using Canu v1.7 (Koren et al., 2017). We polished the Canu contigs with the PacBio long reads using Pbalign v0.3.1 (https://www.pacb.com/support/software-downloads/) and Arrow in the GenomicConsensus package v2.2.2 (https://github.com/PacificBiosciences/GenomicConsensus). We additionally polished the contigs with Illumina (Illumina, San Diego, CA, USA) short reads using Bowtie2 v2.3.5.1 (Langmead and Salzberg, 2012) and Pilon v.1.22 (Walker et al., 2014). The short read-based polishing was iterated until the number of corrected sites did not increase (number of iterations: 18).

Gaps of the short read-based scaffolds (Platanus result) were filled with polished long read-based contigs using an in-house program. The procedure is as follows (Suppl. Fig. 1):

1. Fixed-length regions flanking the gaps in scaffolds were extracted.
2. The flanking sequences were aligned (queried) to the long read-based contigs using Minimap2 v2.17 (Li, 2018) with the options of “-c -k 19.”
3. The alignment results were filtered out if sequence identity <90%, query coverage <25%, or multiple best-scoring hits were identified.
4. Each gap was filled with a region between the alignments in the contig if the following conditions were satisfied:

i. The flanking region pair was aligned to the same contig.
ii. The distance between alignments <50 kbp.
iii. The strands of the alignments were consistent.
iv. There was no ‘N’ in the corresponding regions in the contig.
5. Steps (1) to (4) were iterated eight times. Here, the specific length of the flanking regions was applied for each iteration (500, 1k, 5k, 10k, 20k, 40k, 80k, and 160k).
6. Step (5) was iterated twice.

Finally, to eliminate contamination, we performed homology searches of the assembly against the NCBI genome database (human, bacteria, and virus), the mitochondrial genome of *A. planci* (accession no. NC007788), and the chromosome sequence of the COTS symbiont (COTS27, accession no. AP019861) using BLASTN v2.7.1 (Altschul et al., 1990). The assembled sequences (scaffolds or contigs) were eliminated as contamination if sequence identity ≥90% and query coverage ≥50% (Suppl. Table 1). We assessed the completeness of the draft genome using BUSCO v.4.0.6 with the Metazoa Odb10 dataset (Seppey et al. 2019). K-mer-based (k = 17) estimations of the sizes of the genome and duplicated regions were also conducted using Jellyfish v.2.2.10 (Marçais and Kingsford, 2011) and GenomeScope v.2.0 (Ranallo-Benavidez et al. 2020).

### 2.3 MIG-seq Analysis

MIG-seq is a population genomic method that can easily detect a moderate number of neutral single nucleotide polymorphisms (SNPs) (Suyama and Matsuki, 2015). Previous studies using MIG-seq on marine species successfully discovered species boundaries and genetic structures that were undetectable using traditional genetic markers such as mitochondrial DNA and nuclear ribosomal internal transcribed spacer 2 (Nakabayashi et al. 2019; Takata et al. 2019).

We used eight pairs of multiplex ISSR primers (MIG-seq primer set 1; Suyama and Matsuki, 2015) with the Multiplex PCR Assay Kit Ver. 2 (TaKaRa Bio Inc., Shiga, Japan) for the first PCR; we slightly modified the protocol by Suyama and Matsuki (2015) and used a total reaction volume of 7 μL in a T100™ Thermal Cycler (Bio-Rad, Hercules, CA, USA) under the following conditions: initial denaturation at 94 °C for 1 min; followed by 25 cycles of 94 °C for 30 s, 38 °C for 1 min, and 72 °C for 1 min; and final extension at 72°C for 10 min. The samples were indexed in the second PCR, which was performed with PrimeSTAR GXL DNA polymerase (TaKaRa Bio Inc.) using a total reaction volume of 12 μL in a thermal cycler under the following conditions: 21 cycles of 98 °C for 10 s, 54 °C for 15 s, and 68 °C for 1 min. PCR products were run on an agarose gel, and 350–800 bp PCR products were extracted from the gel. The gel-extracted DNA was pooled and sequenced using MiSeq Control Software v2.0.12 (Illumina) with the MiSeq Reagent v3 150 cycle kit (Illumina). Image analysis and base calling were performed using Real-Time Analysis Software v1.17.21 (Illumina).

To eliminate low-quality reads and primer sequences from the raw data, we used the FASTX-Toolkit v0.0.14 (Gordon and Hannon, 2012; http://hannonlab.cshl.edu/fastx_toolkit/index.html) with a fastq-quality-filter setting of –q 30 –p 40. We removed adapter sequences for the Illmina MiSeq run from both the 5’ end (GTCAGATCGGAAGAGCACACGTCTGAACTCCAGTCAC) and the 3’ end (CAGAGATCGGAAGAGCGTCGTGTAGGGAAAGAC) using Cutadapt v1.13 (Martin, 2011), and then we excluded short reads less than 80 bp. We used Stacks v2.2 (Catchen et al., 2013; Rochette and Catchen, 2017) to stack the reads and extract the SNPs. We used the newly sequenced *A.* cf. *solaris* genome as a reference, as described below. We used the Gstacks option (-rm, -pcr, -duplicates) in Stacks v2.2 to remove PCR duplicates by randomly discarding all but one pair of each set of reads. We used population software in Stacks to prepare the dataset. We used the minimum percentage of individuals required to process a locus across all data (r) equal to 095. After excluding individuals missing more than 10% of SNP data, we used a single SNP option to avoid strong linkage between loci for Structure analysis. We used all SNPs for the remaining genetic analyses.

### 2.4 Population Genetics

BayeScan v2.1 (Narum and Hess, 2011) was used to detect possible SNPs under natural selection with a default setting. Genetic diversity (expected and observed heterozygosities, *H*_*E*_ and *H*_*O*_, respectively), Hardy-Weinberg Equilibrium, global *F_ST_* including and excluding Ogasawara samples, and pairwise *F*_*ST*_ values across different populations were calculated using GenAlEx v6.502 (Peakall and Smouse, 2012) with 9999 permutations.

To test whether the Ogasawara population is genetically different from other populations, a hierarchical analysis of molecular variance (AMOVA) was conducted with prior grouping (Ogasawara vs other populations) using Arlequin v3.5 (Excoffier and Lischer, 2010).

Bayesian clustering analysis was conducted using Structure software v2.3.4 (Prichards et al., 2000) with 500,000 burn-in followed by 500,000 steps of Markov chain Monte Carlo simulations and 10 iterations. We used admixture- and allele frequency-correlated models that were expected to best explain the data set. The LOCPRIOR model (Hubisz et al. 2009) was used with Ogasawara vs other populations and without prior regional grouping, and the number of putative clusters (*K*) was set from 1 to 7. The online software Structure Harvester (Earl and von Holdt, 2012) was used to summarize the likelihood value. Structure plots were summarized using the online software CLUMPAK (Kopelman et al., 2015). To further visualize the genetic relationships among different populations, principal coordinate analysis (PCoA) was conducted using GenAlEx v6.502 based on pairwise genetic distances among populations.

PGDspider v2.0.8.3 (Lischer & Excoffier, 2012) was used to convert the data formats.

### 2.5 Oceanographic larval dispersal simulation

We conducted a Lagrangian larval dispersal simulation to elucidate the oceanographic connectivity between the regions. For this simulation, we used Connectivity Modeling System v2.0 (Paris et al., 2013) with the ocean analysis/reanalysis products produced by the Global Ocean Forecasting System 3.0 of Global HYCOM (Chassignet et al., 2007), which are 1/12° resolution (~9 km) daily products archived from 1993 to 2018 (https://www.hycom.org/). The surface water current and sea surface temperature (SST) in the Global HYCOM products ranged from 115°E–155°E longitude and 15°N– 40°N latitude. Thirty source and sink sites fitting the purpose of this study were chosen to create source sink polygons (see Figure 1A). The particles regarded as COTS larvae were numerically released from ocean areas inside the polygons and tracked. Once the particle entered a polygon, the particle was removed and judged as larval settlement success, and the connection between the source polygon and sink polygon was counted. The particle release timing (start time of COTS mass spawning) and duration of the spawning at each site were determined by the following rules using a history of SST at the site provided by Global HYCOM analysis/reanalysis products based on Yasuda et al. (2010): (1) If SST at the site rose above 28 °C, the particles were released until SST dropped below 26 °C or 30 days passed; (2) If SST at the site did not reach 28 °C, the particles were released while SST was higher than 26 °C. A total of 100 particles per day were released from each source polygon under the above conditions. Finally, we obtained a connectivity matrix that was cleated based on larval dispersal simulations of 26 years (1993 to 2018).

Because the connectivity based on the larval dispersal simulation is a one-generation direct linkage, indicating that stepping-stone migration had not been considered, we conducted back-tracing of the origin site through a discrete-time Markov chain using the connectivity matrix. To this end, we first set the following form of the connectivity matrix **N**(*m*×*m*):

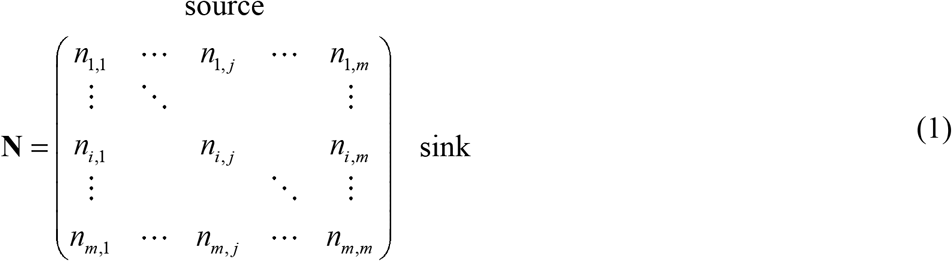

The matrix elements were normalized by the total number of sink particles as follows:

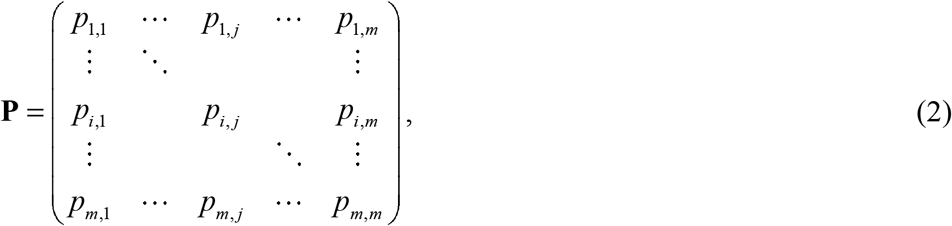

where

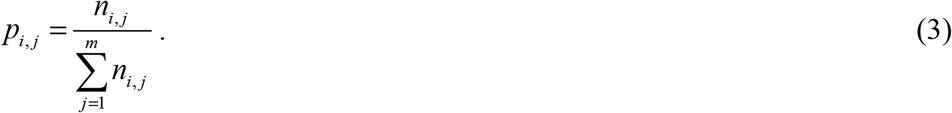

The normalized connectivity matrix can be regarded as the transition matrix of the Markov chain. The *n*-th power of the transition matrix **P**^*n*^ represents the *n* time-step back-trace to the origin source.

## 3 Results

### 3.1 Sequencing and SNP metrics

In total, 11,461,866 raw reads ranging from 9,190 to 281,474 reads per sample were obtained for 187 *A.* cf*. solaris* samples. After filtering out the low-quality reads, 6,125,996 reads ranging from 5,996 to 107,468 reads per sample remained. On average, 189,536 reads per individual remained after two-step filtering. We excluded seven samples missing more than 10% data and used a total of 180 samples for analysis. Using all SNP options, we obtained a total of 464 SNPs, which were used for all subsequent analyses except for Structure analysis. For Structure analysis, 115 SNPs using a single SNP option were used. BayeScan indicated that all loci were neutral (*q*-values >0.05).

### 3.2 Genetic diversity and genetic differentiation

Genetic diversity (*H*_*O*_ and *H*_*E*_) of the studied populations, including that from Ogasawara, were comparable and ranged from 0.039–0.044 and 0.039–0.045 for *H*_*O*_ and *H*_*E*_, respectively (Suppl. Table 2). Global *F*_ST_ values were significant (*P* < 0.001) but small both for groups including (*F*_ST_ = 0.005) and excluding the Ogasawara population (*F*_ST_ = 0.006). Likewise, AMOVA with prior grouping (Ogasawara vs other populations) indicated that *F*_CT_ was low and not significant (0.00167, *P* = 0.219). Pairwise *F*_ST_ values across the studied populations were also small, and only two pairs (Sekisei Lagoon [SKS] and Tatsukushi [TTK], Miyazaki [MYZ] and Ogasawara [OGS]) became significant after sequential Bonferroni correction (α < 0.05) (Table 1).

**Table 1.** Pairwise *F*_*ST*_ values (below) and their probability values (above). Significantly differentiated pairs after sequential Bonferroni correction (α < 0.05) are shown in bold face.

|  | TTK | MYZ | SKR | ONN | MYK | SKS | OGS |
| --- | --- | --- | --- | --- | --- | --- | --- |
| TTK |  | 0.295 | 0.224 | 0.105 | 0.003 | **0.000** | 0.107 |
| MYZ | 0.003 |  | 0.400 | 0.142 | 0.023 | **0.000** | 0.141 |
| SKR | 0.004 | 0.001 |  | 0.465 | 0.484 | 0.21 | 0.473 |
| ONN | 0.006 | 0.002 | 0.000 |  | 0.370 | 0.031 | 0.447 |
| MYK | 0.018 | 0.007 | 0.000 | 0.001 |  | 0.355 | 0.045 |
| SKS | **0.028** | **0.014** | 0.002 | 0.004 | 0.001 |  | 0.004 |
| OGS | 0.006 | 0.003 | 0.000 | 0.000 | 0.005 | 0.008 |  |

Mean log likelihood values (mean lnP(D)) were highest at *K* = 1 without prior grouping and comparable at *K* from 1 to 7 with prior grouping, with the lowest standard deviation value of mean lnP(D) found at *K* = 1 (Suppl. Table 3). Structure bar plots with and without prior geographic grouping suggested genetic homogeneity across the populations (Fig. 2A, Suppl. Fig. 2). PCoA indicated that, along the x-axis, SKS, MYK and Sakurajima [SKR] were divided from the others, although it was not related to geographic location or the year of sampling (Fig. 2B).

**Figure 2.**
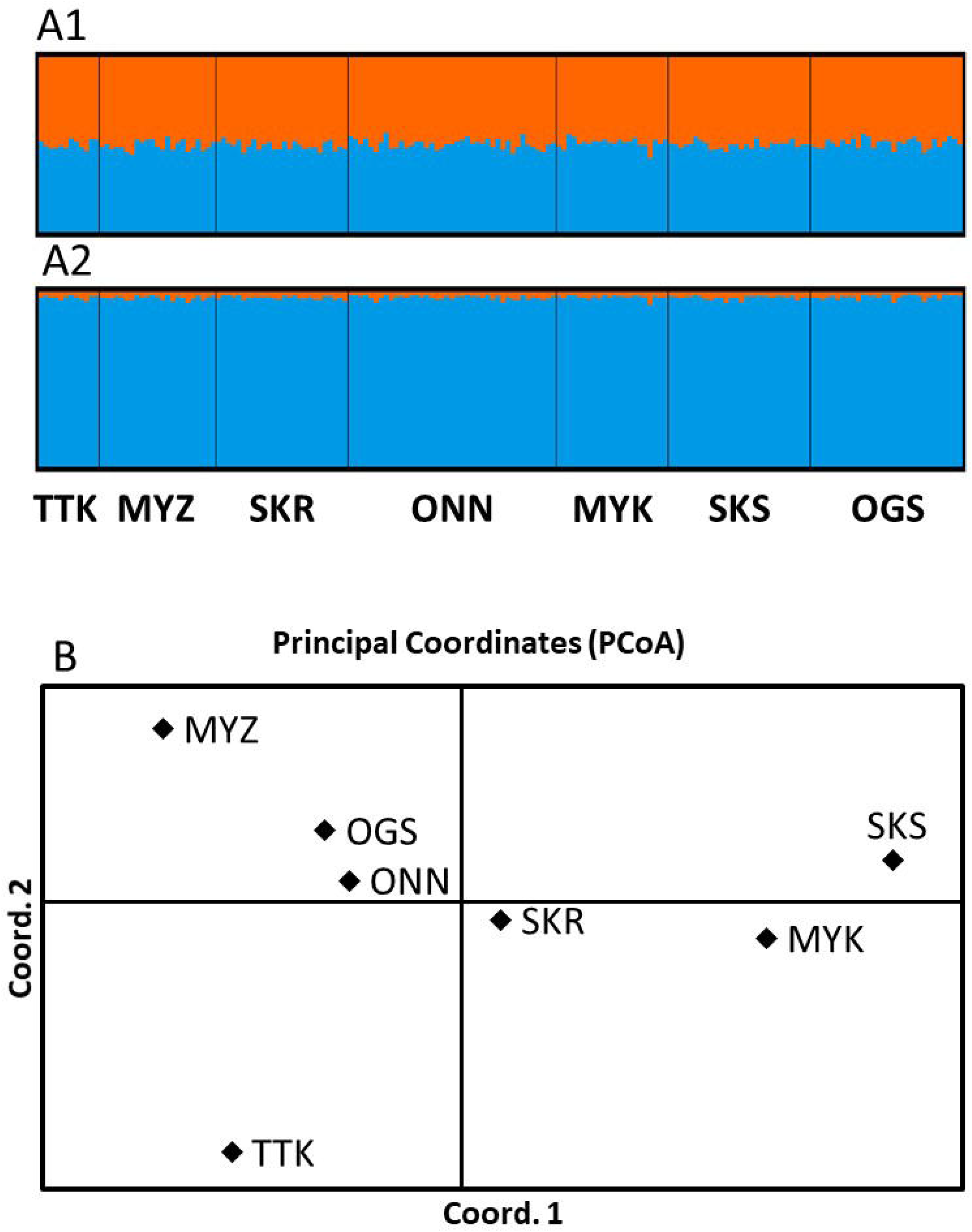
**(A)**: Bar plot from Structure analysis at *K* = 2. **A1**: without prior geographic grouping. **A2**: with prior geographic grouping (Ogasawara vs other regions), indicating the population structure estimated by Structure v2.3.4 (Pritchard et al., 2000). The x-axis indicates individuals, and the y-axis indicates the proportion of hypothetical clusters, shown in different colors. (**B)**: Results of PCoA based on a covariance matrix with standardized data. The first and second axes explain 70.03% and 21.20% of the variation, respectively.

### 3.3 The larval dispersal simulation

The results of the larval dispersal simulations are shown as a connectivity matrix (Fig. 1B). The diagonal elements of the matrix indicate self-seeding. Basically, site numbers from 1 to 20 were assigned from south to north along the Kuroshio Current; site numbers from 21 to 25 were assigned from Nii-Jima and Shikine islands, which are closer to Japan’s main island, to the Ogasawara and Iwo-Jima Islands, which are further; and site numbers from 26 to 30 were assigned along the Tsushima Warm Current. The linkage of the site numbers for numerical simulation to site names for genomic analysis was as follows: 5 = Sekisei Lagoon (SKS), 6 = Miyako (MYK), 7 = Onna Village (ONN), 11 = Miyazaki (MYZ), 13 = Tatsukushi (TTK), 25 = Ogasawara (OGS), and 27 = Sakurajima (SKR). The results of this analysis confirmed that the larvae were transported from south to north by following the Kuroshio Current. However, they also confirmed that self-seeding dominated in the Ogasawara area, and it is very rare for larvae from the other sites to reach the Ogasawara area directly or for larvae from the Ogasawara area to reach the other sites directly.

Figure 3 shows the back-tracing results obtained using Markov chains. The bar of each site indicates the ratio of the number of particles released from the source site (laws of transition matrix **P**^*n*^). The results indicate that the origin sites gradually converged with some of the upstream sites of the Kuroshio Current. The population in the Ogasawara area is not well mixed with that of the Kuroshio region, but the 200–500 time-steps, which may be regarded as representing over a few hundred generations, confirmed that the populations in the Kuroshio region and the Ogasawara region are almost mixed.

**Figure 3.**
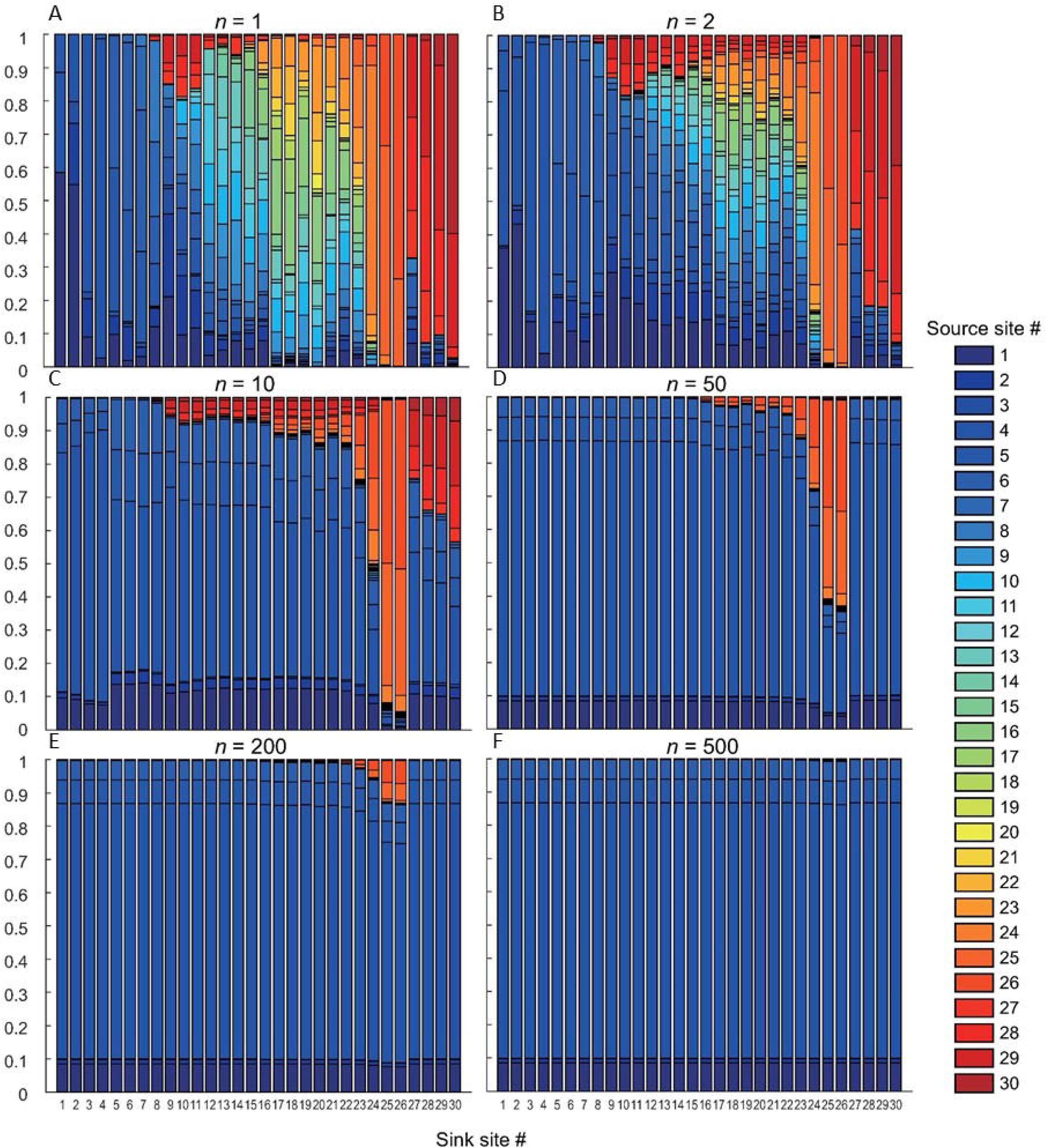
The ratio of the number of particles originating from each source site. **(A)** Simulation of 26-year larval dispersal (see sites, Fig. 1A and connectivity matrix, Fig. 1B). The ratio determined by Markov chains with **(B)** time-step *n* = 2, **(C)** *n* = 10, **(D)** *n* = 50, **(E)** *n* = 200, and **(F)** *n* = 500. Site numbers are assigned as follows: 5 = Sekisei Lagoon (SKS), 6 = Miyako (MYK), 7 = Onna Village (ONN), 11 = Miyazaki (MYZ), 13 = Tatsukushi (TTK), 25 = Ogasawara (OGS), and 27 = Sakurajima (SKR).

## 4 Discussion

This study examined larval dispersal of the coral predator *A.* cf. *solaris* between the Kuroshio region and Ogasawara using population genomic analysis and oceanographic simulation to test the secondary outbreak hypothesis. Contrasting results were obtained using different methods. Population genomic analysis indicated genetic homogeneity between the populations of the two regions, and oceanographic numerical simulation indicated that multiple-generation stepping-stone migration was required to disperse larvae from the Kurshio region to Ogasawara. These results indicate that the Ogasawara population of *A.* cf. *solaris* has the same genetic origin as that of the Kuroshio region, but a large amount of one-generation larval dispersal, such as a secondary outbreak from the Kuroshio region to Ogasawara, is unlikely. Rather, the data suggest that the population outbreak in Ogasawara might have been caused by successful self-seeding. This study highlights the importance of an integrated approach for estimating larval dispersal, including its timescale, for conservation purposes.

### 4.1 Potential of secondary outbreak between Ogasawara and Kuroshio region

In this study, all population genomics results indicated genetic similarity and no genetic structure between *A.* cf. *solaris* populations in the Ogasawara and Kuroshio regions. The AMOVA and Structure results indicate that the origins of *A*. cf. *solaris* populations in the Ogasawara and Kuroshio regions are genetically similar. Low *F*_ST_/*F*_CT_ values between the Ogasawara and Kuroshio regions indicate (1) a contemporary, large amount of larval dispersal, (2) accumulation of stepping-stone migrants over multiple years, or (3) recent separation of a population with no or limited ongoing gene flow. Based on the oceanographic numerical simulation analysis results and historical records of *A*. cf. *solaris* in Ogasawara, the latter two are more likely than contemporary, large amounts of larval dispersal.

The result of the 26-year oceanographic simulation suggests that the chance of direct one-generation larval dispersal between the Kuroshio region and Ogasawara is very low, although stepping-stone migration (e.g. via Hachijyo-jima in Izu islands) between the two regions over multiple generations is physically possible. The Markov chain simulation, which simulated stepping-stone migration over multiple generations, indicated that the Ogasawara population is dominated by migrants from the Kuroshio region after 200–500 generations. This implies that the accumulation of larvae from low levels of migration between the Kuroshio and Ogasawara regions over multiple years would result in genetic homogeneity. Given that there are no observed reports of population outbreaks of *A*. cf. *solaris* in stepping-stone regions (e.g., Izu islands) in the last 40 years (Yasuda 2018), genetic homogeneity between the Ogasawara and Kuroshio regions observed by *F*-statistics and STRUCTURE analysis could be attributed to stepping-stone larval migration over multiple generations. Crandall et al. (2018) demonstrated observed results of the marine species along Hawaii archipelagos where *F*_*ST*_ is not significantly different from zero were not actually panmictic but having weak hierarchical structure of isolation-by-distance caused by stepping stone migration. Coalescent simulation including *A*. cf. *solaris* samples from Izu islands would confirm this hypothesis.

Alternatively, it is possible that the observed strong gene flow resulted from recent colonization. According to a literature survey, there was no record of *A.* cf. *solaris* in the Ogasawara region before 1945, and the first record of this species in Ogasawara was in 1968 (Kurata, 1984; Yasuda, 2018). Since then, the density of *A.* cf. *solaris* in Ogasawara has been low (Yasuda 2018). However, a lack of a record of *A.* cf. *solaris* in Ogasawara before 1945 does not necessarily mean a complete absence of this species in the region, given that almost no studies or surveys were conducted before 1945. In addition, recent migration with a small population size would predict low genetic diversity due to the founder effect, although we observed almost the same genetic diversity in the Ogasawara and Kuroshio regions. Therefore, high genetic diversity in Ogasawara and strong gene flow between the Ogasawara and Kuroshio regions is more likely caused by stepping-stone larval migration over multiple generations, as suggested by the oceanographic simulation.

Whichever may be the case, the oceanographic simulation suggests that contemporary, one-generation larval dispersal between the Ogasawara and Kuroshio regions is limited, and thus direct secondary population outbreaks that require a large amount of larval dispersal within a generation from the Kuroshio region to Ogasawara are unlikely.

Kurata (1984) speculated that *A.* cf. *solaris* in Ogasawara might have colonized either from the Kuroshio region or from Mariana regions such as Guam. According to a population genetics analysis using microsatellite markers, genetic differentiation between the Kuroshio and Guam regions has been identified (Tusso et al., 2016), implying that colonization and/or secondary population outbreaks from Guam to the Ogasawara region are less likely than from the Kuroshio region.

Only one previous study examined the genetic connectivity of coral reef organisms between the Kuroshio and Ogasawara regions. A reef-building coral species, *Acropora* sp., which has a shorter (average 3–4 days) larval stage than that of *A.* cf*. solaris*, showed genetic differentiation between the Ogasawara and Kuroshio regions (Nakajima et al., 2012). This implies that larval dispersal from the Kuroshio region to the Ogasawara region of coral reef organisms with short larval stages is quite limited.

### 4.2 Possible causes of population outbreak in Ogasawara

Oceanographic simulation suggests that many larvae are self-recruited in Ogasawara compared with other *A.* cf. *solaris* populations in the Kuroshio region (Suppl. Table 4). This fact indicates that the observed population outbreak in Ogasawara in 2018 was caused by a sudden increase in regional self-seeding (which is termed “primary outbreak” as opposed to “secondary outbreak”; Birkeland and Lucas, 1990). While several hypotheses have been proposed to explain the primary outbreak of *A.* cf *solaris*, the causes of primary outbreaks are still unclear (Pratchett et al., 2014). While the larval survival hypothesis in association with an elevated level of nutrients is widely accepted in GBR, causes of outbreak would also depend on several regional environmental conditions being met simultaneously (Okaji et al., 2019): elevated levels of larval nutrition and subsequent higher survival rates; lower predation on larvae and juveniles; and physical processes acting on larvae that increase their survival, such as weather conditions and current patterns (Keesing and Halford, 1992; Okaji et al., 2019).

The population outbreak in Ogasawara in 2018 could be attributed to successful local larval recruitment in 2016, given that it takes two years for *A*. cf. *solaris* larvae to become adults (Birkeland and Lucas 1990). Oceanographic simulation also suggests that the number of self-recruited larvae in Ogasawara was larger (397) in 2016, which exceeded the average (252) over 26 years (Suppl. Table 5). This might be partly due to the smaller number of typhoons during the larval period (July to October, estimated by the spawning period associated with water temperature; Yasuda et al., 2010); in 2016, three typhoons passed through the Ogasawara region, which is below the 70-year average (4.47) (data from Japan Meteorological Agency, Suppl. Table 6). A previous field survey discovered dense clouds of *A.* cf. *planci* larvae before one typhoon, but they were scattered and disappeared from the coral reef area after the typhoon (Suzuki et al., 2016), implying that typhoons prevent mass settlement of larvae. However, in some years with a higher number of self-recruits and fewer typhoons, no population outbreak was observed in Ogasawara. Thus, other factors such as higher success of fertilization, higher nutrient levels during the larval stage, and less predation during the post-settlement stage might also be associated with the demographic fluctuation of *A.* cf. *solaris* in the Ogasawara area.

For conservation of corals in the Ogasawara area, caution should be taken two years after the observation of local retention of ocean current systems and preferable conditions for the increase in *A.* cf. *solaris*, such as a large amount of terrestrial run-off and a fewer number of typhoons during spawning periods. Local monitoring of juvenile starfish would be useful for predicting population outbreaks (Okaji et al. 2019).

### 4.3 Multiple approach to estimate larval dispersal in marine organisms

This study estimated larval dispersal of *A.* cf. *solaris* between Kuroshio and Ogasawara using two different approaches: population genomic analysis and larval dispersal simulation. Gene flow estimation by the population genomic approach often reflects both contemporary and evolutionary processes, rendering the interpretation of the amount of contemporary dispersal difficult because of the long evolutionary timescale. Genetic homogeneity can be observed due to a large amount of contemporary larval dispersal (secondary population outbreak), or stepping-stone migrants over multiple generations, or recent divergence with no contemporary dispersal. Our study is unique in that it included an oceanographic model with Markov chain simulation. Such a simulation was useful for testing these alternative hypotheses. A simple oceanographic simulation averaged over 26 years first rejected the secondary population outbreak hypothesis by indicating limited physical larval transport from the Kuroshio to the Ogasawara region within a generation. This result supports the idea that stepping-stone migrants over multiple generations is responsible for the genetic homogeneity observed in the *F*_*ST*_ and STRUCTUTE analysis between the Kuroshio and Ogasawara populations.

This may give the impression that oceanographic simulation alone is enough for testing the secondary population hypothesis; however, it is generally imperfect for estimating larval dispersal. Our model regarded larvae as neutral passive particles without any of the biological characteristics of larvae that may influence actual larval dispersal, such as vertical movement, natural selection, food availability, and mortality. In addition, the model only simulates the dispersal process, which does not include post-settlement survival. In some marine species, genetic connectivity is quite limited despite having long pelagic larval duration possibly caused by several biological features of larvae and juveniles (e.g. Barber et al. 2000; Barber et al. 2002). Thus, a comparison with genetic connectivity that reflects all the biological characteristics of the species is also important. While several previous studies have reported good agreement between oceanographic models and genetic analysis and have provided robust and straightforward interpretations of larval dispersal occurring on an ecological timescale (e.g. Galindo et al. 2006; White et al. 2010; Sunday et al. 2014; Taninaka et al. 2019), only a few have discussed the usefulness of oceanographic modeling for identifying alternative hypotheses or provided additional insights into the causes of observed population genetic structure (Galindo et al. 2010; Crandall et al. 2014). By applying Markov chain simulation in the numerical simulation, we could also include the effect of migration over multiple generations, which is easier to compare with genetic results than a single generation simulation result in terms of time-scale. This method underlines the usefulness and importance of using both genetic and oceanographic methods to estimate larval dispersal and provides significant insight into larval dispersal that occurs on ecological and evolutionary timescales.

## 5 Author Contributions

NY conceived the study. MH, NY, RK, HY, and TY drafted the manuscript. RK, HY, TI, and AT sequenced, assembled, and reconstructed the reference genome. NY, YA, and TS collected the *Acanthaser* samples. MH, HT, HY, TK, and NY conducted the MIG-seq analysis. TN and TY conducted oceanographic simulations. All authors edited the draft and approved it for publication.

## 7 Acknowledgments

We are grateful for Ms. Eriko Nagahiro for her help in the molecular experiment.

## Data Availability Statement

The whole genome sequencing data (BioSample ID, SAMD00056692) and MIG-seq data (BioSample: SSUB016768, SAMD00282440-SAMD00282626) were submitted to DDBJ and available.

## 12. Supplementary Captions

**Suppl. Fig. 1** Schematic model of gap-filling with long read-based contigs. Flanking-region lengths: 500, 1k, 5k, 10k, 20k, 40 kbp, 80 kbp, 160 kbp (multiple values are applied). Gap-closing step is iterated two times.

**Suppl. Table 1.** Statistics of the draft genome.

**Suppl. Table 2** Sample information and genetic diversity inbreeding coefficient. N = number of individuals analyzed. *Ho* = observed heterozygosity, *He* = expected heterozygosity, *F* = inbreeding coefficient.

**Suppl. Table 3.** The results from Structure Harvester to calculate *ΔK*. *K* is the number of hypothetical clusters assumed in Structure analysis.

**Suppl. Figure 2. Structure analysis at comparable *K* from 3 to 7** (1 = Tatsukushi [TTK], 2 = Miyazaki [MYZ], 3 = Sakurajima [SKR], 4 = Onna Village [ONN], 5 = Miyako [MYK], 6 = Sekisei Lagoon [SKS], 7 = Ogasawara [OGS]). **(A)**: without prior and **(B)**: with prior grouping for Ogasawara. In K = 6, each pattern was supported in five out of ten iterations. The x-axis indicates sampling sites. The y-axis indicates the probability of membership.

**Suppl. Table 4. The total number of arrived particles over 26 years.** Bold face numbers indicate self-seeding. Self-seeding at Ogasawara is highlighted in yellow. Vertical columns indicate source populations, and horizontal rows indicate sink sites.

**Suppl. Table 5.** The average number of arrived particles over 26 years. Bold face numbers indicate self-seeding. Self-seeding at Ogasawara is highlighted in yellow. Vertical columns indicate source populations, and horizontal rows indicate sink sites.

**Suppl. Table 6.** The number of typhoons approaching Ogasawara and Izu Islands over a 70-year span (1951–2020).

