## Supplemental Materials for "Integrated population genomic analysis and numerical simulation to estimate larval dispersal of *Acanthaster* cf. *solaris* between Ogasawara and other Japanese regions"

Supplementary Material

### 1 Supplementary Figures and Tables

### Supplementary Table 1. Statistics of the draft genome

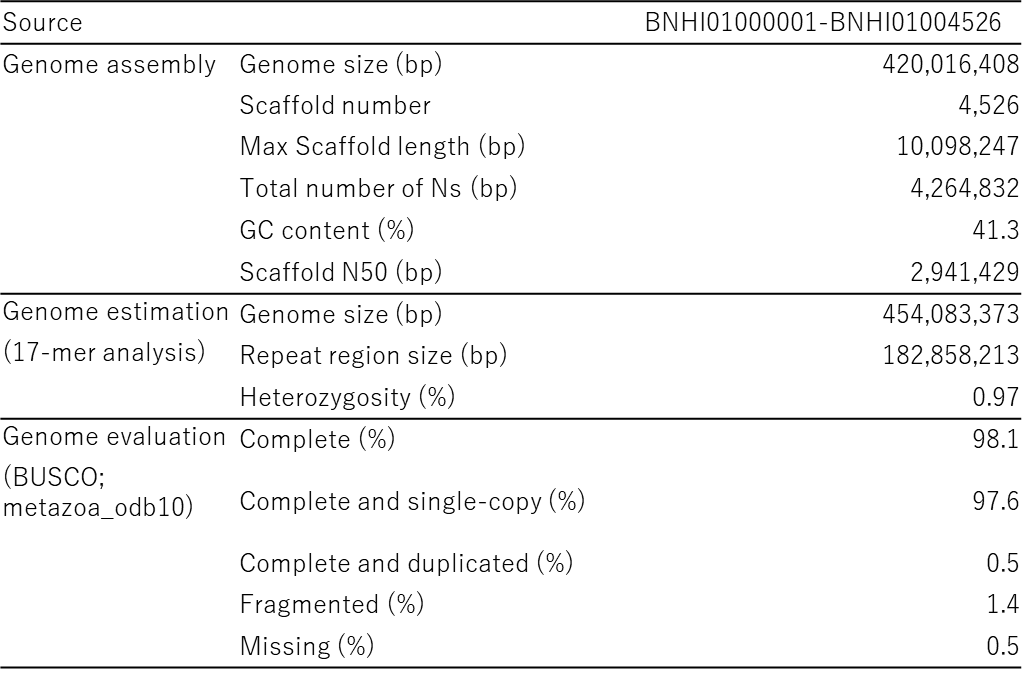

**Supplementary Table 2. Sample information and genetic diversity inbreeding coefficient**

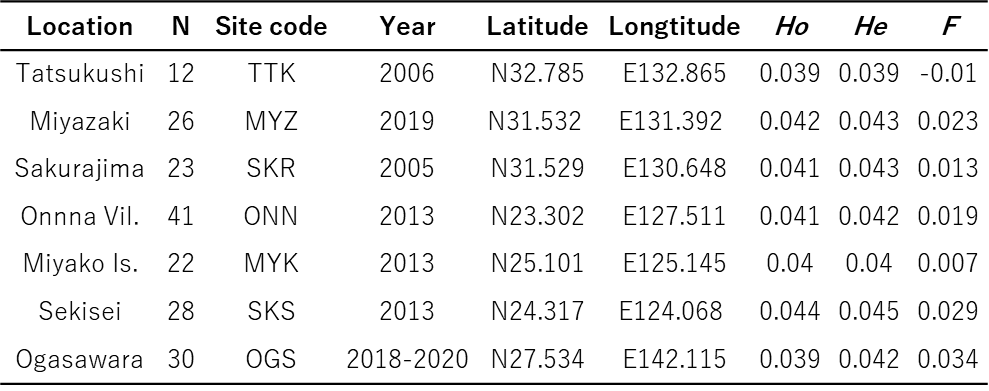

N = number of individuals analyzed. *Ho* = observed heterozygosity, *He =* expected heterozygosity, *F* = inbreeding coefficient.

### Supplementary Table 3. The results from Structure Harvester to calculate ΔK

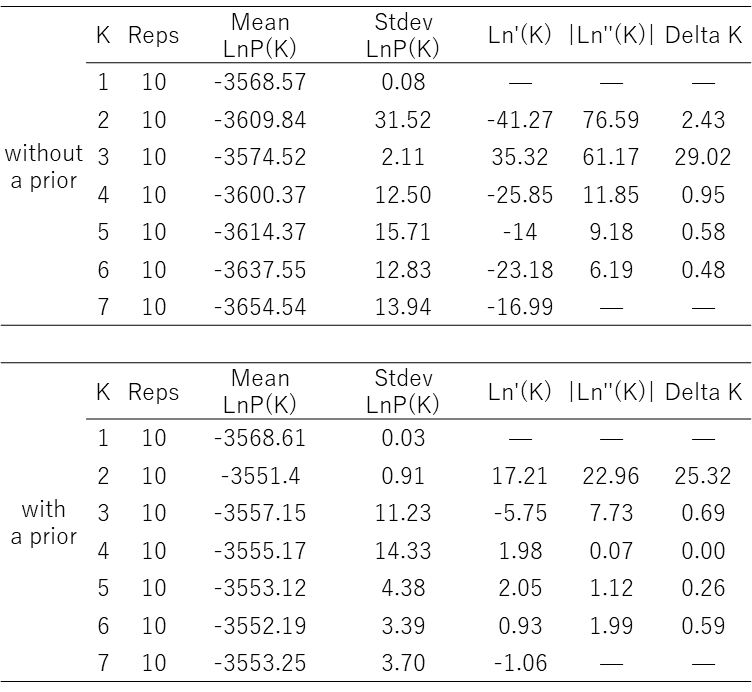

K is the number of hypothetical clusters assumed in Structure analysis.

**Supplementary Table 4. The total number of arrived particles over 26 years**

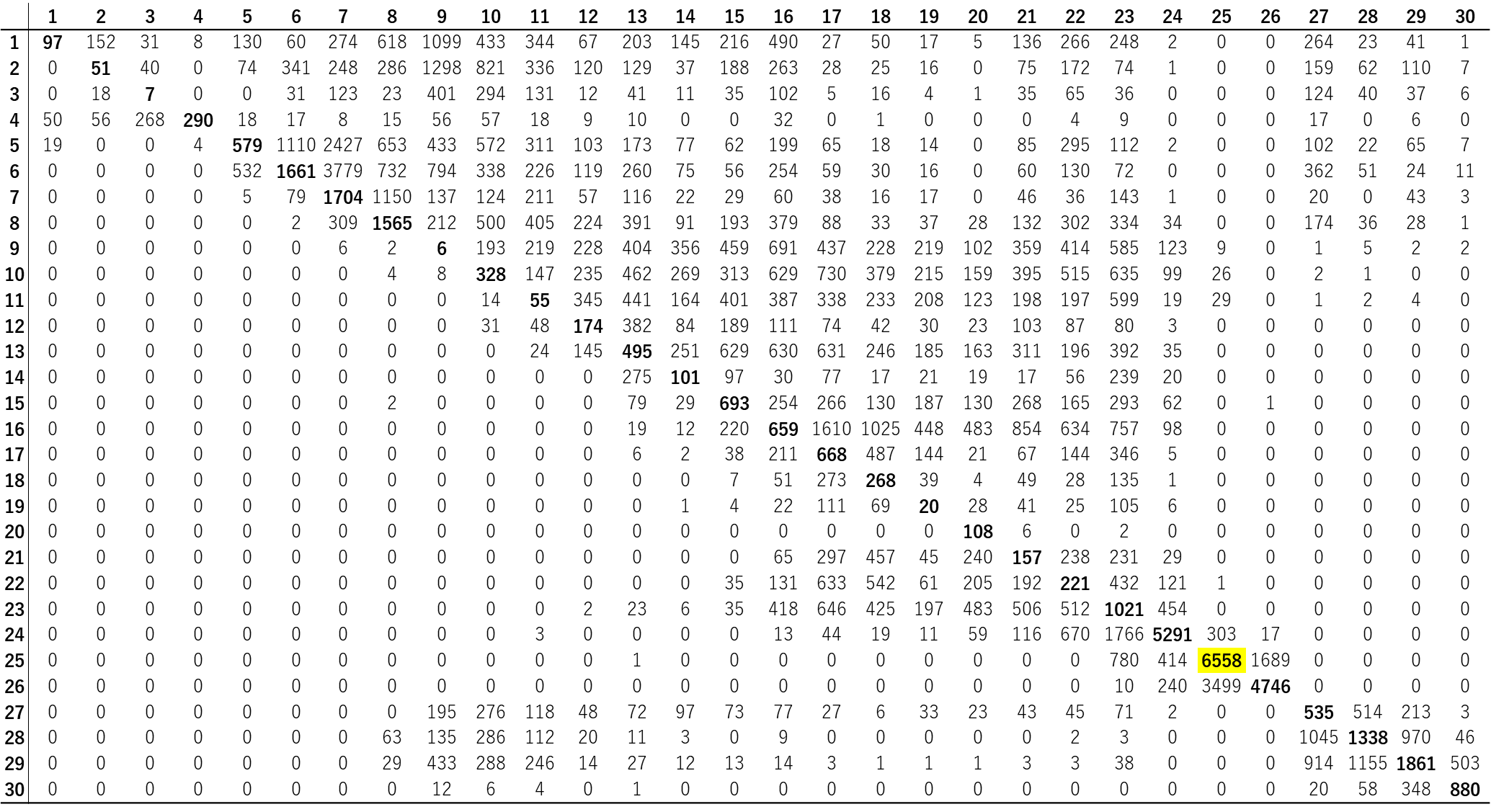

Bold face numbers indicate self-seeding. Self-seeding at Ogasawara is highlighted in yellow. Vertical columns indicate source populations, and horizontal rows indicate sink sites.

### Supplementary Table 5. The average number of arrived particles over 26 years

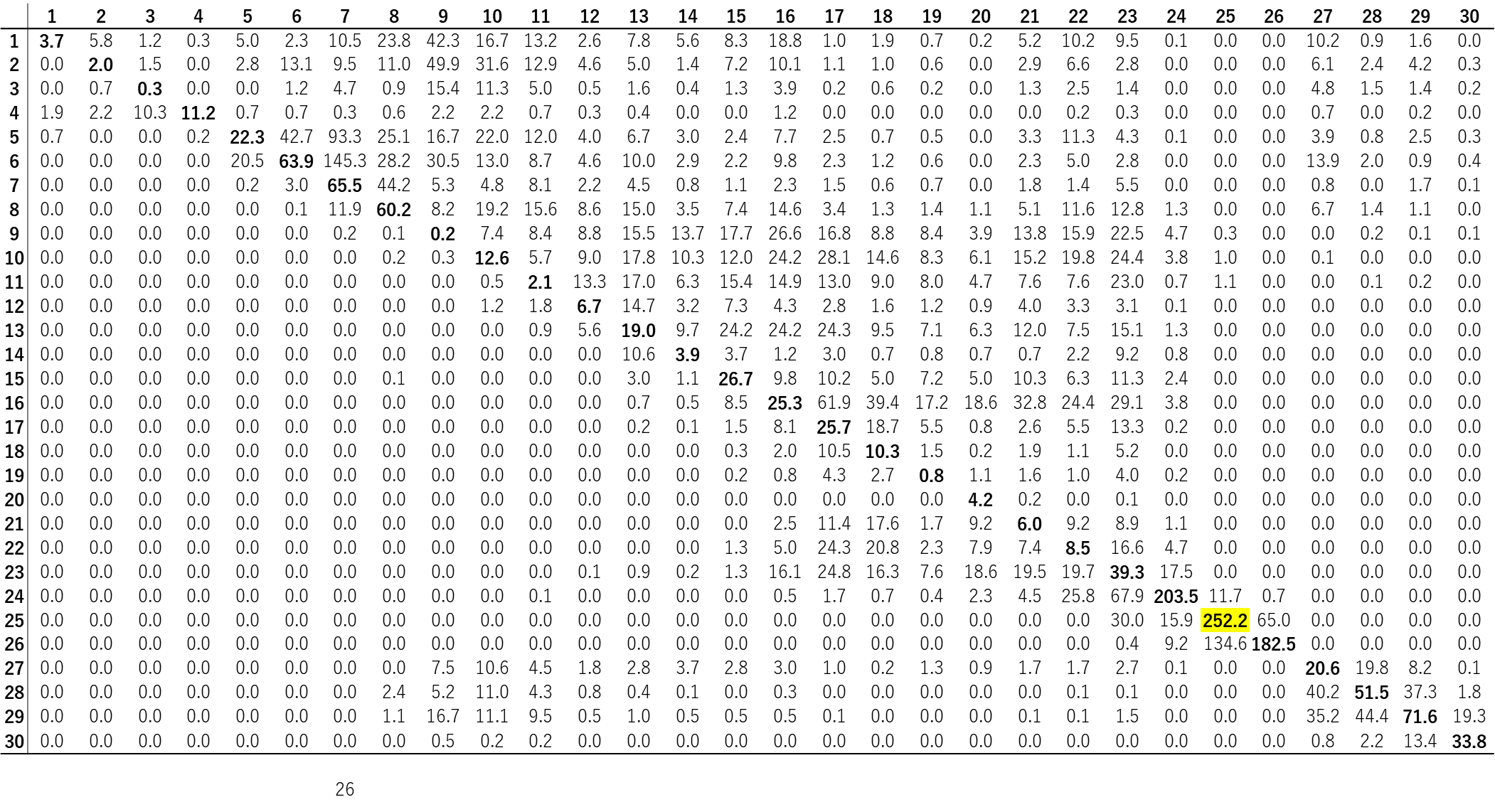

### Bold face numbers indicate self-seeding. Self-seeding at Ogasawara is highlighted in yellow. Vertical columns indicate source populations, and horizontal rows indicate sink sites.

**Supplementary Table 6. The number of typhoons approaching Ogasawara and Izu Islands over a 70-year span (1951–2020)**

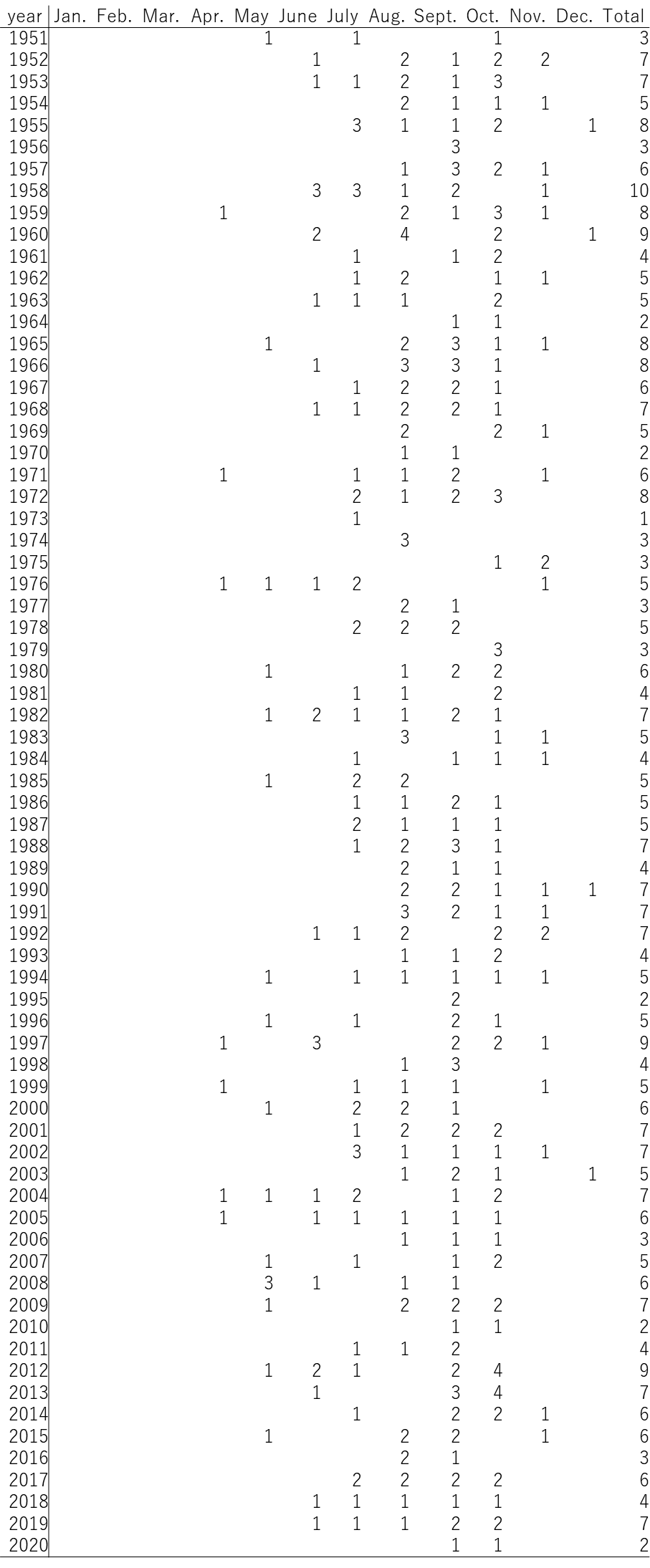

For more information on Supplementary Material and for details on the different file types accepted, please see [here](http://home.frontiersin.org/about/author-guidelines" \l "SupplementaryMaterial). Figures, tables, and images will be published under a Creative Commons CC-BY licence and permission must be obtained for use of copyrighted material from other sources (including re-published/adapted/modified/partial figures and images from the internet). It is the responsibility of the authors to acquire the licenses, to follow any citation instructions requested by third-party rights holders, and cover any supplementary charges.

#### 1.1 Supplementary Figures

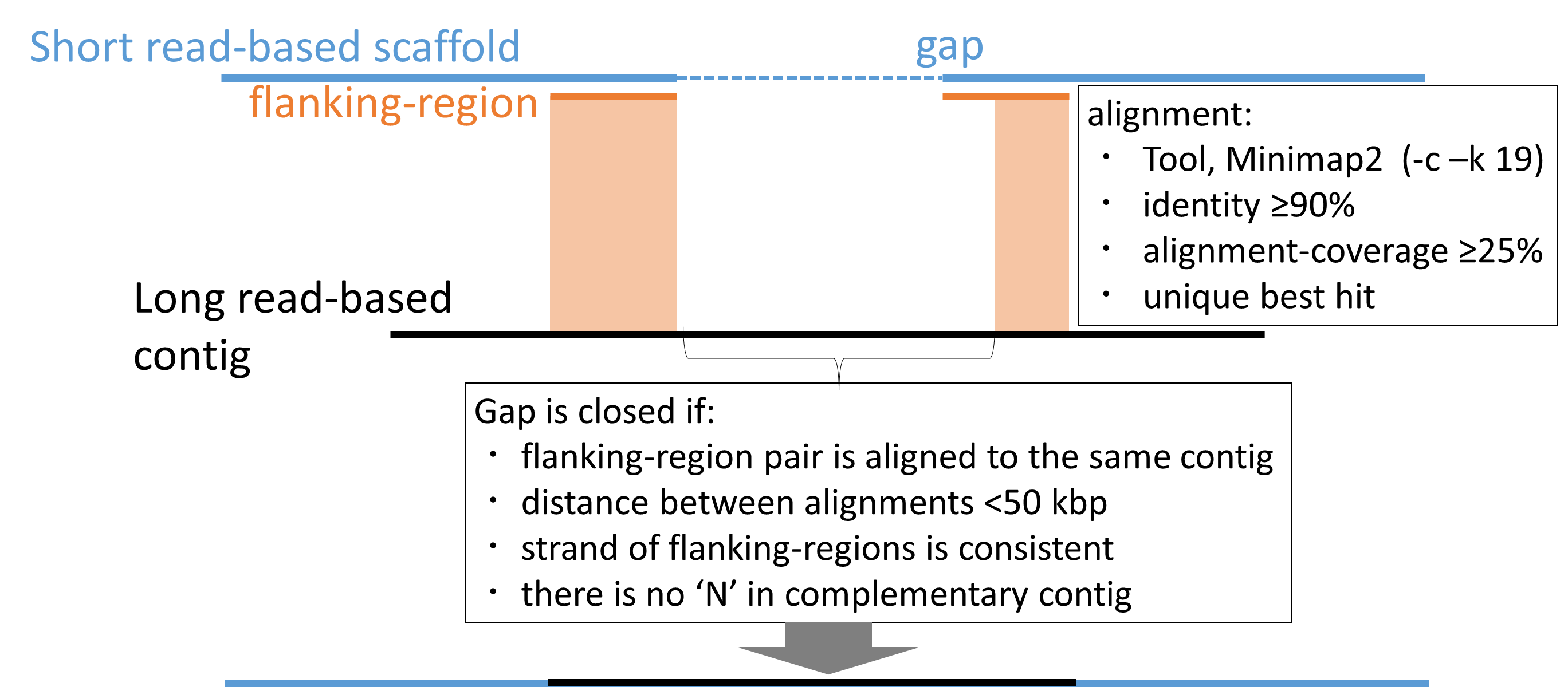

**Supplementary Figure 1.** Schematic model of gap-filling with long read-based contigs. Flanking-region lengths: 500, 1k, 5k, 10k, 20k, 40 kbp, 80 kbp, 160 kbp (multiple values are applied). Gap-closing step is iterated two times.

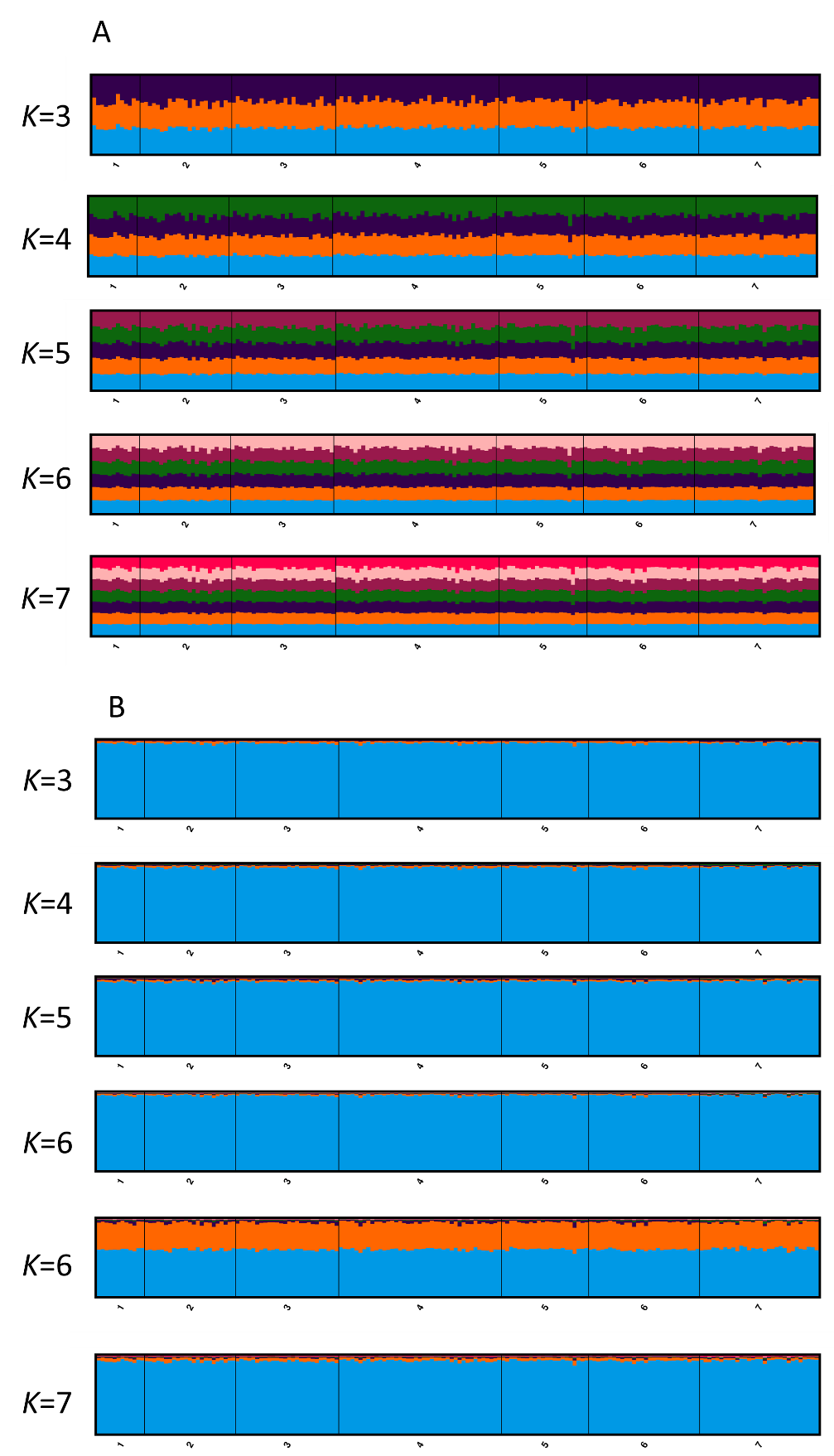

**Supplementary Figure 2.** Structure analysis at comparable *K* from 3 to 7 (1 = Tatsukushi [TTK], 2 = Miyako [MYO], 3 = Sakurajima [SKR], 4 = Onna Village [ONN], 5 = Miyazaki [MYK], 6 = Sekisei Lagoon [SKS], 7 = Ogasawara [OGS]), (A): without prior and (B): with prior grouping for Ogasawara. In K = 6, each pattern was supported in five out of ten iterations. The x-axis indicates sampling sites. The y-axis indicates the probability of membership.
